# Epi-PoreC: a nanopore-based method for simultaneous profiling of chromatin conformation, DNA methylation, and chromatin accessibility

**DOI:** 10.64898/2026.09.15.751767

**Authors:** Anastasia McKinlay, Wei Zong, Craig S. Pikaard

## Abstract

Understanding how the three-dimensional (3D) organization of the genome relates to chromatin and epigenetic landscapes requires methods capable of measuring chromosomal DNA interactions and chromatin modifications in an integrated manner. Current approaches typically use separate assays to assess chromatin conformation and epigenetic state, thus these features are not assessed on the same DNA molecules. Here, we describe Epi-PoreC, an Oxford Nanopore long-read sequencing-based method that combines the chromosome conformation capture method, PoreC with Fiber-seq, a method for detecting accessible chromatin regions based on their ability to be labeled using an exogenous adenosine methyltransferase. The positions of chromosome contacts, 6-methyladenosines, and endogenous 5-methylcytosines are then detected by nanopore sequencing, all on the same DNA molecules. Using *Arabidopsis thaliana* nuclei, we demonstrate that Epi-PoreC yields genome-wide chromatin contact maps comparable to those obtained by standard PoreC and DNA methylation and accessibility profiles comparable to standard Fiber-seq experiments. Epi-PoreC is thus an efficient means for conducting epigenetic and 3D genome profiling on individual DNA sequences in a single assay.

**Multidisciplinary abstract:** Understanding genome regulation is aided by integrating information about chromosome architecture, DNA methylation, and chromatin accessibility. However, most experimental methods measure these features separately, precluding their detection on the same DNA molecules.

Here we describe Epi-PoreC, an nanopore sequencing–based method that simultaneously measures chromatin contacts, DNA methylation, and chromatin accessibility in a single assay. The approach integrates chromosome conformation capture (PoreC) with Fiber-seq–based adenine methyltransferase labeling of accessible DNA. Using *Arabidopsis thaliana* nuclei, we demonstrate that Epi-PoreC produces chromatin contact maps highly concordant with those obtained by standard PoreC assays and DNA methylation and accessibility profiles comparable to those obtained by Fiber-seq.

Because long nanopore sequencing reads can traverse multi-way chromatin ligation products, Epi-PoreC enables detection of chromatin interactions and modifications on the same DNA molecules, yielding genome folding and epigenetic landscape information relevant to studies of gene and genome function. Epi-PoreC therefore expands the toolkit for multi-omic singlemolecule genome analysis, particularly for complex or repetitive genomic loci where long-read sequencing is needed to identify sub-repeats or sequence elements that map to specific chromosomal locations.

**Method summary:** Epi-PoreC integrates Fiber-seq–based adenine methyltransferase labeling of accessible DNA with the PoreC chromosome conformation capture protocol. Long-read Oxford Nanopore sequencing then allows the simultaneously detection of chromatin contacts, cytosine methylation and chromatin accessibility on individual DNA molecules. The method thus provides an efficient workflow for integrated single-molecule analyses of genome architecture and epigenetic state.

## Introduction

The functional organization of the genome is reflected, in part, by the interplay between higherorder chromatin organization and epigenetic state (1,2). Chromosome conformation capture methods can reveal how short and long-range chromatin interactions contribute to 3D genome folding (3,4) and sequencing-based methods for mapping DNA methylation or chromatin accessibility can illuminate features of the epigenetic landscape (5–7).

Long-read Oxford Nanopore sequencing technology has enabled the development of methods to extract multiple layers of information from individual DNA molecules (5–7) and is invaluable for the study of repetitive regions where only long-read sequencing can identify patterns of subtle variation unique to specific sets of sub-repeats (8–12). The PoreC method uses nanopore sequencing of nuclear DNA that has been crosslinked, digested into fragments, and then ligated in order to identify junctions between non-contiguous segments of chromosomal DNA (13). The method can reveal multi-way chromatin contacts and high-resolution maps of 3D genome organization (14). The Fiber-seq method uses an N6-adenosine methyltransferase to methylate the subset of nuclear adenosines accessible to the enzyme (7,15,16). Nuclear DNA is then purified and subjected to Nanopore sequencing and basecalling, allowing methylated and unmethylated adenosines to be differentiated. Basecalling also allows differentiation of unmodified cytosines from cytosine methylated by endogenous cytosine methyltransferases. Fiber-seq thus provides simultaneous measurement of relative DNA accessibility and endogenous cytosine methylation at single nucleotide resolution.

Here, we describe Epi-PoreC, a method that integrates Fiber-seq with PoreC in a single workflow (Figure 1A), enabling chromosome contact mapping, chromatin accessibility and cytosine methylation analyses on single DNA molecules. We show that Epi-PoreC yields results that are comparable to the PoreC (13) and Fiber-seq (7) methods performed separately.

**Figure 1.**
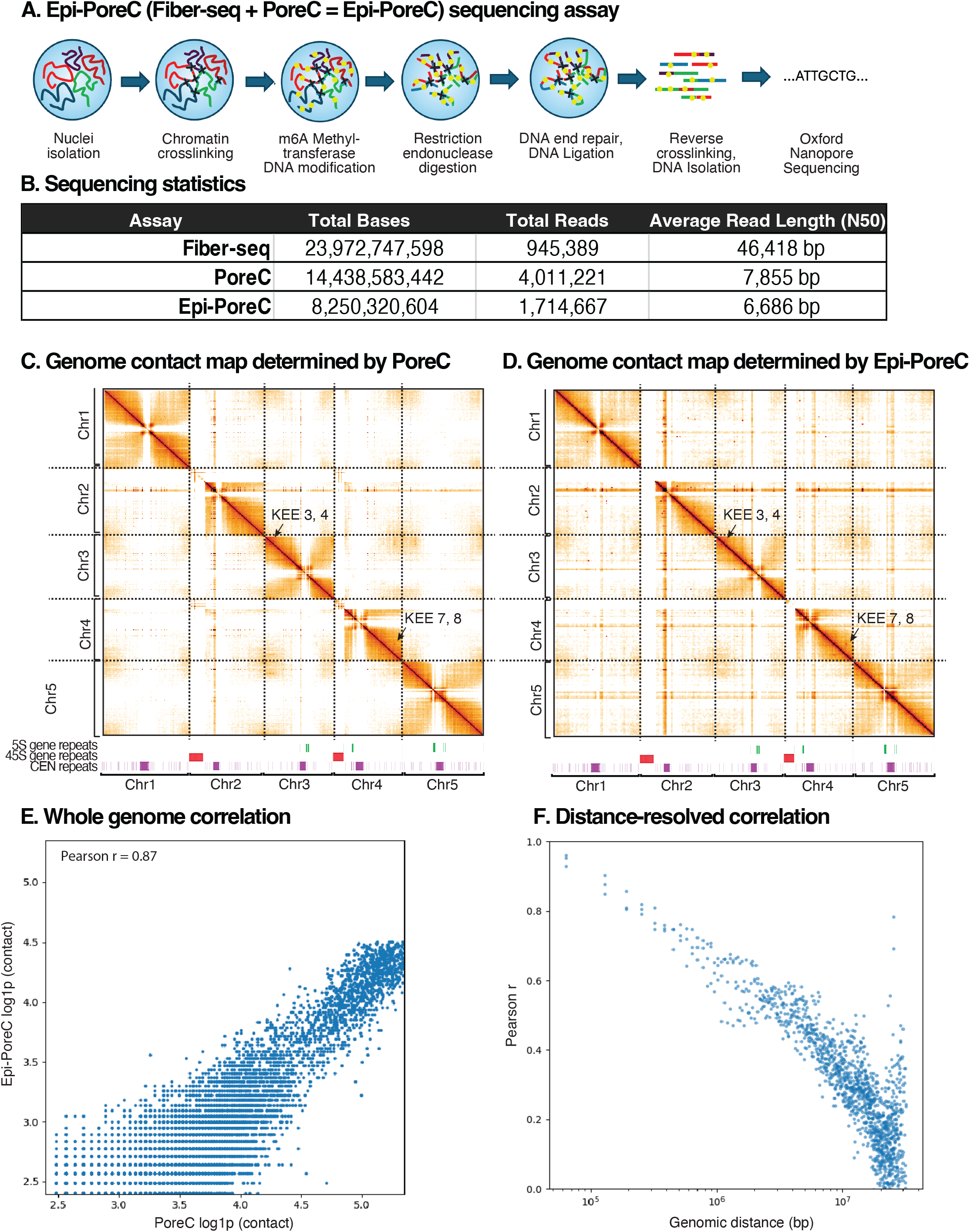
Overview and testing of the Epi-PoreC assay. **(A)** Schematic of the Epi-PoreC workflow that integrates adenosine methylation labeling with PoreC chromatin conformation capture. Intact nuclei are isolated and subjected to chemical crosslinking to fix chromatin interactions. Accessible chromatin regions are preferentially labeled using an adenine methyltransferase, after which chromatin is digested with a restriction enzyme and subjected to ligation to join DNA ends in proximity to one another. Following reversal of crosslinks and DNA purification, long DNA fragments containing multi-way ligation products are sequenced using Oxford Nanopore Technology. **(B)** Sequencing statistics for Fiber-seq, PoreC (with HindIII digestion), and Epi-PoreC (with HindIII digestion) libraries, including total bases sequenced, total read counts, and read length distributions (N50). **(C)** Genome-wide chromatin contact map generated using standard PoreC, displayed for all five *thaliana* chromosomes. Prominent regions, including nucleolus organizer regions (*NOR2* and *NOR4*), centromeric repeats, and KNOT Engaged Elements (KEE3/4 and KEE7/8)(19), are indicated. **(D)** Genome-wide chromatin contact map generated using Epi-PoreC. Large-scale interaction patterns and known chromatin features are similar to those obtained using the standard PoreC protocol (see panel **C**). **(E)** Whole-genome comparison of chromatin contact frequencies obtained by PoreC or Epi-PoreC. Contact matrices were generated at 64-kb resolution and filtered to retain bin pairs with a minimum of 10 raw contacts. Contact counts were log1p-transformed prior to comparison. Genome-wide contact frequencies derived from Epi-PoreC show strong concordance with those obtained from standard PoreC (Pearson r = 0.87). **(F)** Distance-resolved correlation analysis of chromatin contact frequencies between PoreC and Epi-PoreC. Intra-chromosomal contact pairs were stratified by genomic separation into logspaced distance bins, and Pearson correlation coefficients were calculated within each bin after trimming extreme values (2nd–98th percentiles). High correlation is observed at short and intermediate genomic distances, with a gradual decrease at larger separations, consistent with expected distance-dependent contact sparsity.

## Materials and Methods

### Plant material

*Arabidopsis thaliana* accession Columbia-0 (Col-0; Arabidopsis Biological Resource Center stock CS1092) plants were grown under long-day conditions (16 h light / 8 h dark). Unopened inflorescence apices were harvested, flash frozen in liquid nitrogen, and stored at −80 °C until use.

### Nuclei isolation and chromatin crosslinking

Nuclei were isolated from approximately 600 mg of frozen inflorescence tissue according to the PacBio TissueRuptor Plant Tissue Protocol (NUC-TRP-001). Tissue was homogenized in a 50 mL conical (Falcon) tube in 20 mL Nuclei Isolation Buffer (NIB) prepared according to the PacBio/Circulomics TissueRuptor Plant Tissue Protocol (NUC-TRP-001) and supplemented with β-mercaptoethanol (50 µL of 14 M β-mercaptoethanol per 20 mL NIB). The homogenate was then incubated on a rotating mixer (20 rpm) at room temperature for 20 min. Chromatin crosslinking was performed by adding formaldehyde to a final concentration of 1% and incubating at room temperature for 15 min with rotation. Crosslinking was quenched by addition of glycine to a final concentration of 0.125 M and incubation for 5 min. Nuclei were filtered through 20 µm vacuum filters (Steriflip) in 50 mL conical tubes and pelleted by centrifugation in a swinging bucket rotor at 7,000 × g for 20 min at 4 °C. Pellets were washed by gentle resuspension in 20 mL NIB followed by centrifugation as above. Three washes were conducted. Following the final wash, nuclei were resuspended in 1 mL ice-cold phosphate-buffered saline (PBS; 1× PBS: 137 mM NaCl, 2.7 mM KCl, 10 mM Na□HPO□, 1.8 mM KH□PO□; pH 7.4), transferred to 1.5 mL Protein LoBind microcentrifuge tubes (Eppendorf), and pelleted at 1,000 × g for 5 min at 4 °C.

### PoreC and Epi-PoreC library preparation

PoreC libraries were prepared following the Oxford Nanopore Technologies Plant PoreC protocol (https://nanoporetech.com/document/extraction-method/plant-pore-c) with minor modifications. Crosslinked nuclei pellets in 1.5 mL Protein LoBind microcentrifuge tubes (Eppendorf) were resuspended in 100 µL 0.5% SDS prepared in nuclease-free water and incubated at 62°C for 6 min, followed by cooling to room temperature. Next, 250 µL of nuclease-free water and 50 µL of 10% v/v Triton X-100 were added to the nuclei suspension, mixed gently by pipetting with a wide-bore pipette tip (Thermo Scientific™ ART™, Cat # 21-236-2C), and incubated at 37 °C for 15 min without agitation. Restriction endonuclease digestion was performed using HindIII in 1x CutSmart buffer (New England Biolabs), using 500 U enzyme per reaction and incubation at 37 °C for 18 h without agitation. Heat inactivation of the nuclease was performed at 80 °C for 20 min prior to proximity ligation. T4 DNA ligase proximity ligation, proteinase K digestion, crosslink reversal, and DNA purification were carried out as described in the Oxford Nanopore protocol (https://nanoporetech.com/document/extraction-method/plant-pore-c).

For Epi-PoreC, steps of the Fiber-seq protocol were added to the PoreC protocol prior to HindIII restriction digestion. First, the nuclei permeabilization step was performed with 0.05% digitonin in Wash Buffer on ice for 15 min. Wash Buffer consisted of 20 mM HEPES-KOH (pH 7.5), 150 mM NaCl, 0.5 mM spermidine, 0.1% BSA, and one EDTA-free Roche Complete protease inhibitor tablet per 50 mL buffer. After washing the nuclei pellet twice with 0.95 mL Tween-Wash buffer (Wash Buffer supplemented with 0.1% (v/v) Tween-20), nuclei were transferred to 1.5 mL Protein Low-Bind tubes (Eppendorf). Nuclei were then subjected to the activation step, in which the nuclei pellet was gently resuspended in 100 μL Activation Buffer (15 mM Tris-HCl (pH 8.0), 15 mM NaCl, 60 mM KCl, 1 mM EDTA, 0.5 mM EGTA, 0.05 mM spermidine, and 0.1% BSA, with S-adenosyl methionine added to a final concentration of 800 μM immediately before use). 200 nM pA-Hia5 (a protein A-Hia5 adenine methyltransferase fusion protein) was then added to catalyze N6-methyladenine (6mA) modification of accessible adenosines. The reaction was performed at 37°C for 2 hr with occasional gentle mixing performed by repeated pipetting with a wide-bore tip (Thermo Scientific™ ART™, Cat # 21-236-2C) every 30 min. S-adenosyl methionine was replenished after 1 hour of incubation. Upon completion of the reaction, the sample was subjected to centrifugation at 7,000 × g for 5 min at 4°C (Eppendorf centrifuge 5424 R). Pellets were then resuspended and subjected to restriction digestion, proximity ligation, and library preparation as described above.

Nanopore sequencing libraries (~100 ng DNA) were prepared using the Oxford Nanopore Rapid DNA Sequencing Kit (SQK-RAD114) and sequenced using PromethION R10 flow cells for 72 hrs. Rapid sequencing chemistry was selected for Epi-PoreC libraries to accommodate reduced input material following proximity ligation.

### Protein A-Hia5 fusion protein expression and purification

Plasmid pD861-SR-pA-Hia5 (kindly provided by the Altemose lab), encoding a His-tagged Protein A-Hia5 fusion protein, was transformed into Escherichia coli BL21-AI™ (Invitrogen™, C607003) cells. One colony was picked and grown overnight in 20 mL LB media at 37°C to obtain a starter culture. To induce protein expression, 20 mL starter culture was added to 1 L LB and grown at 37°C on a rotating shaker until the cell density reached an optical density at 600 nm of 0.6. Then 0.1 mM L-rhamnose was added to induce protein expression, and the culture was grown for another 3 hours at 37 °C. Subsequently, the cells were collected by centrifuging at 7,000 g for 10 min in 500 mL Beckman centrifuge tubes, washed once with phosphate-buffered saline (PBS; 1× PBS: 137 mM NaCl, 2.7 mM KCl, 10 mM Na□HPO□, 1.8 mM KHLPOL; Ph 7.4), flash frozen in liquid nitrogen and stored at −80 °C. To purify the protein, the pellet was resuspended in 30 mL lysis buffer (25 mM HEPES, pH 7.6, 400 mM NaCl, 10% v/v glycerol, 0.1% IGEPAL, 1X Bugbuster® (Sigma, 70584), 20 mM imidazole, 0.5 mM DTT, 1 mM PMSF, 1X Protease Inhibitor cocktail (Roche)) and transferred to 50 mL Oak Ridge Centrifuge tube, PPCO (Thermo Scientific, Cat # 3119-0050). 3 µL/mL Lysonase (Sigma, 71230) was added to the lysis buffer and incubated on a rotating mixer at 4 °C for 10 min to lyse the cells. After centrifuging for 30 min at 50,000 g (fixed angle rotor, Beckman Coulter Avanti J-26 XPI), the supernatant was transferred to a 50 mL Falcon tube with 2 mL Ni-NTA Agarose (Qiagen) for 1 hour at 4 °C with gentle mixing by rotation. The Ni-NTA agarose was collected by centrifuging at 250 g at 4 °C for 5 min (Eppendorf centrifuge 5910 R), transferred into an empty chromatography column, and washed with 20 mL wash buffer (25 mM HEPES, pH 7.6, 400 mM NaCl, 10% v/v glycerol, 0.1% IGEPAL, 40 mM imidazole, 0.5 mM DTT, 0.1 mM PMSF). Bound protein was eluted with 6 mL elution buffer (25 mM HEPES, pH 7.6, 400 mM NaCl, 5% v/v glycerol, 250 mM imidazole, 0.5 mM DTT). Next, the eluted protein was concentrated using a Pierce™ Protein Concentrator (10K MKCO) to 500 µL and further purified by FPLC size exclusion chromatography using a SuperDex 200 Increase 10/300 GL column (Cytiva) equilibrated and run in 20 mM HEPES, pH 7.6, 100 mM NaCl, 5% v/v glycerol column buffer. The purified protein sample was concentrated using a Pierce™ Protein Concentrator (10K MKCO) to a final volume of 400 µL, aliquoted at 10 µL per microcentrifuge tube, and stored at −80 °C.

### Fiber-seq (open chromatin–directed DNA methylation)

Fiber-seq libraries were generated from isolated nuclei as described by Maslan et al. (Fiberseq/DiMeLo-seq) (16). Nuclei were resuspended in 1 mL Wash Buffer (20 mM HEPES-KOH pH 7.5, 150 mM NaCl, 0.5 mM spermidine, 0.1% BSA, and one EDTA-free Roche Complete protease inhibitor tablet per 50 mL buffer), transferred to a 1.5 mL DNA LoBind tube (Eppendorf), permeabilized by addition of digitonin to a final concentration of 0.05% (v/v) and incubated on ice for 15 min. Nuclei were pelleted by centrifugation and washed twice by gentle resuspension and centrifugation in 0.95 mL Tween-Wash buffer (Wash Buffer supplemented with 0.1% Tween-20). For methyladenosine labeling, nuclei were transferred to 1.5 mL Protein LoBind tube (Eppendorf) and gently resuspended in 100 μL Activation Buffer (15 mM Tris-HCl pH 8.0, 15 mM NaCl, 60 mM KCl, 1 mM EDTA, 0.5 mM EGTA, 0.05 mM spermidine, and 0.1% BSA) containing 800 μM S-adenosyl-L-methionine (SAM) and 200 nM pA-Hia5, adenine methyltransferase as described previously. The reaction was incubated at 37°C for 2 hours, with gentle mixing (by hand) every 30 min and replenishment of SAM after 1 hour. Following labeling, nuclei were pelleted by centrifugation at 7,000 × g for 5 min at 4°C, flash-frozen in liquid nitrogen, and stored at −80 °C in 1.5 mL Protein LoBind tube (Eppendorf). Frozen nuclei were subsequently resuspended by 20 sec of vortexing on max speed in 45 μL of Proteinase K, and high-molecular-weight DNA was extracted using the Nanobind HMW Plant Nuclei DNA Extraction Kit (PacBio) according to the manufacturer’s protocol. Libraries (~10 μg DNA) were prepared using the Oxford Nanopore Ultra-Long DNA Sequencing Kit (SQK-ULK114) and sequenced on PromethION R10 flow cells for up to 72 hr.

### Data processing and analysis

Nanopore sequencing data were basecalled using Oxford Nanopore basecalling software with modified base detection enabled. Reads were aligned to the *Arabidopsis thaliana* TAIR12 reference genome using minimap2 (17), with parameters optimized for long-read alignment.

For Pore-C and Epi-PoreC datasets, multi-way ligation products were deconstructed into pairwise chromatin contacts and aggregated into genome-wide contact matrices using the Oxford Nanopore Pore-C analysis workflow (https://nanoporetech.com/document/epi2me-workflows/wf-pore-c). Contact matrices were generated in Cooler (.cool) format (18) and visualized using ResGen (https://github.com/knightlab-analyses/resgen). Prior to comparison, contact counts were filtered to retain bin pairs with a minimum of 10 raw contacts, followed by log1p transformation to reduce the influence of extreme values and enable linear comparison across a broad dynamic range.

Genome-wide chromatin contact concordance between PoreC and Epi-PoreC was assessed by computing Pearson correlation coefficients between log-transformed contact frequencies across all intra-chromosomal bin pairs.

For distance-resolved analyses, intra-chromosomal contact pairs were stratified by genomic separation, calculated as the linear distance between interacting bins. Correlation coefficients were computed within log-spaced distance bins, and extreme values were trimmed by excluding the lower 2nd percentile and upper 98th percentile of contact frequencies to reduce the influence of outliers. Distance-resolved Pearson correlation coefficients were calculated separately for each chromosome and visualized as a function of genomic distance.

DNA cytosine methylation (mCG) and adenine methylation (mA) frequencies were extracted from nanopore signal-level data. Chromatin accessibility was inferred from adenine methylation frequency. Genome-wide comparisons between Fiber-seq and Epi-PoreC results were performed using Pearson and Spearman correlation analyses.

## Results & Discussion

### Sequencing performance of Epi-PoreC

To test whether the Epi-PoreC protocol yields data equivalent to the Fiber-Seq and PoreC assays performed separately, we conducted all three assays in parallel. Sequencing metrics for the Fiber-seq, PoreC, and Epi-PoreC libraries are summarized in Figure 1B. Fiber-seq libraries generated the longest reads, with an N50 exceeding 46 kb, likely due to the absence of restriction endonuclease digestion and proximity ligation steps. Standard PoreC libraries yielded a larger number of reads, but with shorter read lengths (N50 ≈ 7.9 kb), consistent with fragmentation by restriction endonuclease digestion. Epi-PoreC libraries produced fewer total reads and total nucleotides sequenced compared to PoreC libraries, with a modest reduction in read length (N50 ≈ 6.7 kb). This reduction is consistent with the additional enzymatic processing steps required for adenosine labeling and subsequent library preparation. Despite this size decrease, read lengths were sufficient to support detection of multi-way chromatin contacts and extraction of epigenetic landscape information from individual Nanopore reads.

### Genome-wide chromatin contact maps are preserved in Epi-PoreC

Genome-wide chromatin contact maps generated using Epi-PoreC closely resembled those obtained with standard PoreC (Figure 1C–D). Both methods yielded similar interaction patterns for all five *Arabidopsis thaliana* chromosomes, including strong interchromosomal interaction enrichment along the diagonal and prominent long-range interaction features. Distinct interaction signals associated with CEN159 and CEN178, two major centromeric satellite repeat families, were readily observed in both datasets. In addition, previously described higher-order chromatin interaction features, including KNOT Engaged Elements (KEE3/4 and KEE7/8) (19), were detected in both the PoreC and Epi-PoreC contact maps. Together, these observations indicate that chromatin interaction architecture is not compromised by integrating the methyladenosine labeling step of Fiber-seq into the PoreC workflow.

### Quantitative assessment of chromatin contact frequencies detected by PoreC and Epi-PoreC

To quantitatively compare PoreC and Epi-PoreC chromosome contact maps, multi-way ligation products were deconstructed into pairwise contacts and aggregated into genome-wide contact matrices at 64-kb resolution. Contact matrices were filtered to retain intra-chromosomal bin pairs with a minimum of 10 raw contacts and were log1p-transformed prior to comparison. Genomewide comparison revealed strong concordance between the PoreC and Epi-PoreC contact frequencies (Pearson r = 0.87; Figure 1E), indicating that global chromatin interaction patterns are not altered appreciably by the extra adenosine methylation step included only in the Epi-PoreC method.

To further test whether results of the two experimental protocols are in agreement across genomic scales, distance-resolved correlation analysis was performed by stratifying intrachromosomal contact pairs according to genomic separation using log-spaced distance bins. Pearson correlation coefficients were calculated within each bin after trimming extreme values (2nd–98th percentiles). High correlation was observed at short and intermediate genomic distances, with a gradual decrease at larger separations (Figure 1F). This distance-dependent decay is consistent with expected reductions in contact frequency and increased sparsity at long genomic distances and does not indicate systematic disagreement between the datasets.

### DNA methylation profiles generated by Epi-PoreC match Fiber-seq

To evaluate the accuracy of DNA methylation measurements obtained using Epi-PoreC versus Fiber-seq, cytosine methylation frequencies determined by the two methods were compared. Genome-wide comparison of CG methylation frequencies showed extremely high agreement between the two methods (Pearson r = 1.00; Spearman r = 0.93; Figure 2A), demonstrating that the DNA crosslinking, digestion, and ligation steps of the Epi-PoreC protocol do not negatively affect the detection of endogenous 5-methylcytosine positions.

**Figure 2.**
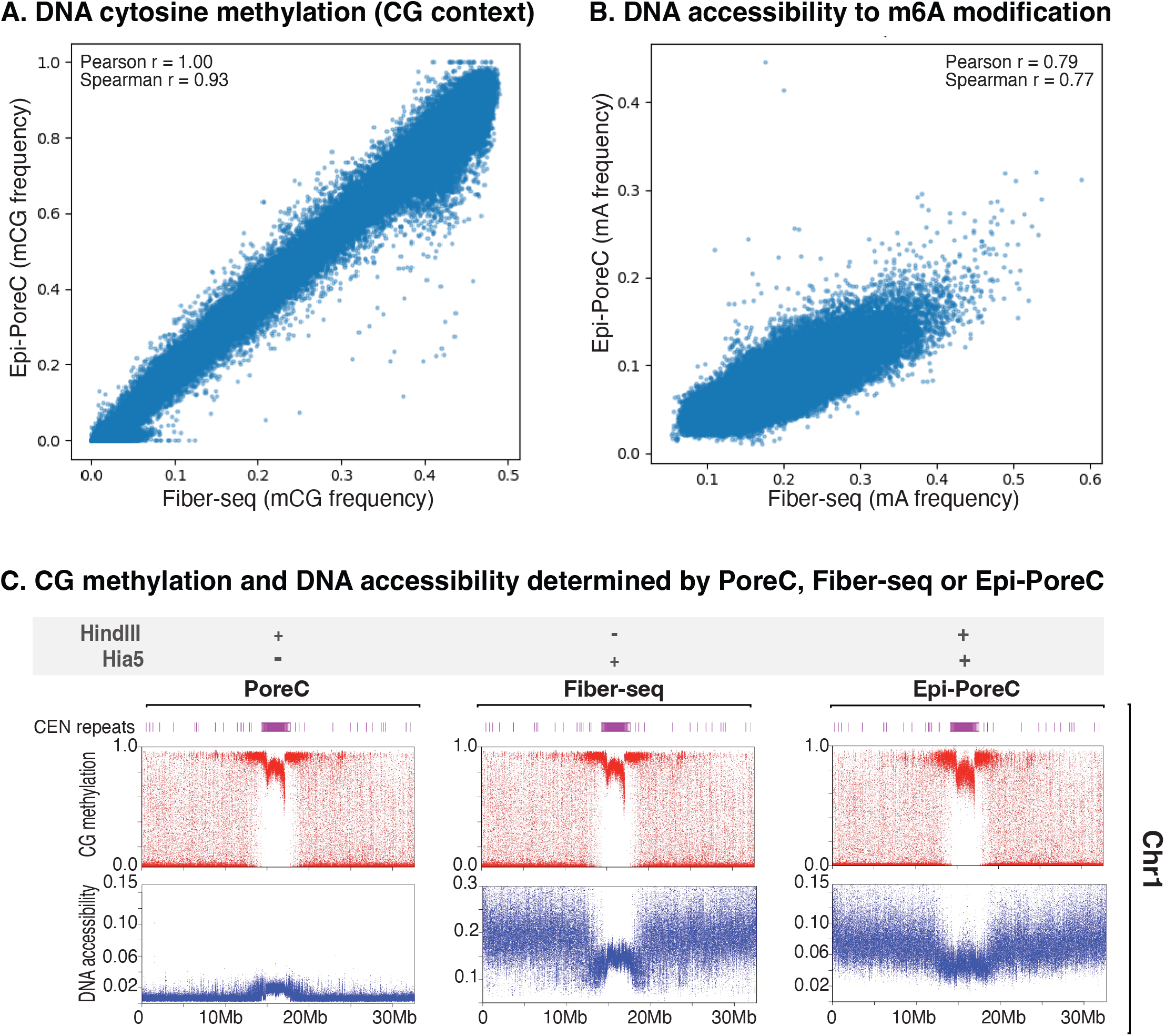
Validation of epigenetic measurements using Epi-PoreC. Comparison of DNA methylcytosine frequencies within CG mptifs detected by Fiber-seq or Epi-PoreC across the genome. High concordance is observed (Pearson r = 1.00; Spearman r = 0.93) for the two methods. Comparison of chromatin accessibility measurements inferred from adenosine methylation frequency determined using Fiber-seq or Epi-PoreC. Accessibility measurements using the two assays show strong correlation (Pearson r = 0.79; Spearman r = 0.77). Chromosome-level profiles of CG DNA methylation and chromatin accessibility across chromosome 1. Tracks reveal similar epigenetic landscapes detected by PoreC, Fiber-seq, or Epi-PoreC, including reduced accessibility and increased cytosine methylation within centromeric regions and higher accessibility and reduced cytosine methylation across chromosome arms.

Chromosome-level visualization further showed that Epi-PoreC recapitulated the methylation patterns observed by Fiber-seq, including elevated CG methylation within centromeric regions and lower methylation levels in the chromosome arms (Figure 2C).

### Chromatin accessibility measured by Epi-PoreC and Fiber-seq

We compared chromatin accessibility detection for the Epi-PoreC and Fiber-seq methods, measured as adenosine methylation frequency throughout the genome. Accessibility measurements showed strong correlation between the two protocols, again suggesting that the DNA crosslinking, digesting, and ligation steps of the Epi-PoreC method do not preclude the detection of 6-methyladenosine (6-mA) positions (Pearson r = 0.79; Spearman r = 0.77; Figure 2B). The 6-mA accessibility measurements did exhibit greater dispersion than the CG methylation measurements, but the overall trends were similar using the two methods.

At the chromosome scale, Epi-PoreC accessibility profiles closely matched those generated by Fiber-seq, with reduced accessibility observed in heterochromatic centromeric repeat regions and higher accessibility across more euchromatic chromosome arms (Figure 2C). These results indicate that the chromosome conformation capture steps of the Epi-PoreC protocol do not substantially compromise chromatin accessibility measurements.

## Conclusions

Our study describes Epi-PoreC, a method that integrates chromosome conformation capture with epigenetic profiling using long-read Nanopore sequencing. Benchmarking analyses demonstrate that Epi-PoreC preserves the genome-wide chromatin contact information measured by PoreC while also generating DNA methylation and chromatin accessibility profiles comparable to Fiber-seq. By combining these techniques within a single experimental protocol, Epi-PoreC enables direct measurements of chromatin interactions and local epigenetic state on the same DNA molecules.

Epi-PoreC offers several advantages over individual assays, including the ability to obtain multiple layers of genomic information from a single experiment, thus reducing the cumulative time and resources required to conduct separate assays. This integration streamlines experimental design and minimizes sample handling while maintaining molecule-level information. The longread sequencing inherent to the approach is particularly advantageous for analyzing repetitive or structurally complex genomic regions that are difficult, or impossible, to resolve using short-read technologies. However, several limitations of Epi-PoreC should be considered. Sequencing read length (N50) is reduced relative to standard PoreC, reflecting the additional enzymatic steps and increased library complexity introduced by adenosine labeling. This reduction may limit resolution in highly repetitive regions that depend on ultra-long reads for accurate mapping of individual repeats. In addition, adenine methylation–based accessibility measurements are an indirect proxy for chromatin accessibility and may be influenced by local sequence context or enzymatic efficiency.

## Future perspective

Advances in long-read sequencing technologies enable the integration of multiple molecular measurements from single DNA molecules. Methods, such as Epi-PoreC, will facilitate studies of how chromatin architecture interacts with epigenetic state to regulate gene expression and genome stability. Future improvements in sequencing throughput, read length, and base-modification detection are likely to further increase the resolution and sensitivity of integrated assays.

Epi-PoreC has the potential to be applied across diverse organisms and cell types to investigate genome organization in development and disease. Coupling the method with targeted protein– DNA interaction labeling or CRISPR-based enrichment strategies may further extend its application to the study of specific genomic loci or regulatory complexes. Together, these developments will contribute to a more comprehensive understanding of how genome structure and epigenetic mechanisms cooperate to regulate genome function.

## Article highlights

- Epi-PoreC integrates chromosome conformation capture with epigenetic profiling in a single nanopore sequencing workflow.
- The method simultaneously detects chromatin contacts, DNA methylation, and chromatin accessibility on individual DNA molecules.
- Genome-wide chromatin contact maps generated by Epi-PoreC show strong concordance with standard PoreC experiments.
- DNA methylation and accessibility measurements are comparable to those obtained using Fiber-seq.
- Epi-PoreC enables integrated single-molecule analysis of genome architecture and epigenetic state.

**Reference annotations**

1. **Sexton et al., 2012** Demonstrated principles of three-dimensional genome organization in *Drosophila* using chromosome conformation capture approach.
2. **Lieberman-Aiden et al., 2009** Introduced the Hi-C method for genome-wide mapping of chromatin interactions, providing the first comprehensive view of 3D genome organization.
3. **Rao et al., 2014** Generated high-resolution Hi-C maps revealing chromatin loops and structural domains that organize mammalian genomes.
4. **Dekker et al., 2013** Review describing chromosome conformation capture technologies and their applications in studying genome architecture.
5. **Simpson et al., 2017** Demonstrated detection of DNA cytosine methylation using nanopore sequencing signals, enabling direct epigenetic profiling with long-read sequencing.
6. **Rand et al., 2017** Described methods for mapping DNA methylation using nanopore sequencing, establishing approaches for detecting epigenetic modifications from raw sequencing signals.
7. **Stergachis et al., 2020** Introduced chromatin fiber sequencing (Fiber-seq), a long-read method for mapping chromatin accessibility and regulatory architecture at single-molecule resolution.
8. **Miga, 2020** Discussed advances in centromere sequencing enabled by long-read technologies and telomereto-telomere genome assemblies.
9. **Naish et al., 2021** Characterized the genetic and epigenetic landscape of *Arabidopsis* centromeres using long-read sequencing and epigenomic profiling.
10. **Fultz et al., 2023** Analyzed sequence and epigenetic differences between active and silent nucleolus organizer regions in *Arabidopsis*, highlighting regulation of repetitive rRNA gene arrays.
11. **Nurk et al., 2022** Reported the first complete telomere-to-telomere assembly of a human genome, demonstrating the power of long-read sequencing to resolve repetitive regions.
12. **Jain et al., 2018** Demonstrated nanopore sequencing and assembly using ultra-long reads, enabling improved characterization of complex genomic regions.
13. **Ulahannan et al., 2019** Introduced the Pore-C method, which uses nanopore sequencing of DNA concatemers to detect multi-way chromatin interactions.
14. **Li et al., 2022** Applied the Pore-C method in *Arabidopsis* to simultaneously capture genome-wide chromatin interactions and DNA methylation information.
15. **Altemose et al., 2022** Introduced DiMeLo-seq, a long-read sequencing method for mapping protein–DNA interactions genome-wide.
16. **Maslan et al., 2024** Provided a detailed experimental protocol for DiMeLo-seq, describing practical implementation of long-read protein–DNA interaction mapping.
17. **Li, H. 2018** Introduced Minimap2, a fast and accurate sequence aligner that has become a standard tool for mapping long- and short-read sequencing data.
18. **Abdennur, N. and L.A. Mirny. 2020** Introduced the Cooler file format, enabling efficient storage, analysis, and visualization of Hi-C contact matrices.
19. **Grob, S., M.W. Schmid, and U. Grossniklaus. 2014**

Provided one of the first high-resolution characterizations of three-dimensional genome organization in Arabidopsis, identifying the KNOT chromatin structure and its long-range interactions.

## Author contributions

A.M. designed and performed experiments, conducted data analysis, and drafted the manuscript.

W.Z. conceived the idea, purified adenosine methyltransferase pA-Hia5 protein, and contributed to experimental interpretation.

C.S.P. supervised the project, contributed to experimental interpretation, and revised the manuscript.

All authors reviewed and approved the final manuscript.

## Data and code availability

Sequencing data have been deposited in the NCBI Sequence Read Archive (SRA) under BioProject PRJNA1495192 and will be publicly released upon publication.

